# Amygdala controls saccade and gaze physically, motivationally, and socially

**DOI:** 10.1101/608703

**Authors:** Kazutaka Maeda, Ken-ichi Inoue, Jun Kunimatsu, Masahiko Takada, Okihide Hikosaka

## Abstract

The amygdala is uniquely sensitive to emotional events. However, it is not understood whether and how the amygdala uses such emotional signals to control behavior, especially eye movements. We therefore injected muscimol (GABA_A_ agonist) into the central nucleus of amygdala (CeA) in monkeys. This unilateral temporary inactivation suppressed saccades to contralateral but not ipsilateral targets, resulting in longer latencies, hypometric amplitudes, and slower velocity. During free viewing of movies, gaze was distributed mostly in the ipsilateral hemifield. Moreover, CeA inactivation disrupted the tendency of gaze toward social interaction images, which were normally focused on continuously. Conversely, optogenetic stimulation of CeA facilitated saccades to the contralateral side. These findings suggest that CeA controls spatially selective gaze and attention in emotional contexts, and provide a new framework for understanding psychiatric disorders related to amygdala dysfunction.

**Highlights:**

- Central amygdala facilitates contralateral saccades selectively.
- Saccade facilitation is related to motivational goals and social interaction.
- The amygdala thus controls goal-directed behavior based on emotional contexts.

## Introduction

Eye movements are important for scanning the visual environment and making decisions. Many psychiatric disorders (e.g., attention-deficit/hyperactivity disorder, bipolar disorder, and autism) are commonly featured by abnormal patterns of eye movements (Munoz et al., 2003; Constantino et al., 2017; Yep et al., 2018). Amygdala dysfunction is thought to be a key cause in these disorders (Davis and Whalen, 2001; Avino et al., 2018). While the amygdala is known to mediate threat detection and passive fear responses, it remains unclear if this structure also contributes to controlling eye movements. Some studies have shown that amygdala lesions alter gaze patterns, especially for face images (Dal Monte et al., 2015; Taubert et al., 2018). Other studies have shown that amygdala neurons are spatially selective and encode information about both the location and the motivational significance of visual cues (Peck and Salzman, 2014). Spatial attention is tightly coupled to motor function, especially in the case of eye movements elicited by stimulus-driven orienting. However, causal evidence linking the amygdala to abnormal eye movement patterns is lacking in both human and animal studies.

We have recently shown that amygdala neurons, mostly in the central nucleus (CeA), encode emotional contexts (dangerous vs. safe and rich vs. poor) (Maeda et al., 2018). Importantly, we found that although neuronal activity differed across subjects, responses of amygdala neurons were correlated with the saccadic reaction time to reward-associated objects within subjects. Anatomical studies have reported that CeA sends output to the basal ganglia, including the caudate tail (CDt), the globus pallidus externus (GPe), and the substantia nigra pars reticulata (SNr), all of which are known to encode stable object values and control saccades through the SNr-superior colliculus (SC) pathway (Shinonaga et al., 1992; Vankova et al., 1992; Fudge and Haber, 2000; Griggs et al., 2017; Amita et al., 2018). These studies suggest that CeA neurons modulate saccadic eye movements by sending contextual information to the basal ganglia circuit.

Here we demonstrate that saccadic eye movements are strongly modulated by muscimol-induced inactivation and optogenetic stimulation of CeA on one side (Supplementary schematic). Our results suggest that CeA controls saccadic eye movements and gaze position in a highly lateralized manner.

## Results

### CeA inactivation suppressed contralateral saccade

We first locally inactivated CeA by injecting the GABA_A_ receptor agonist, muscimol (8.8 or 44 nmol in a 1µl volume) while monkeys performed a visually-guided saccade task (Fig. 1a) to examine whether CeA modulates saccadic eye movements and gaze positions.

**Fig. 1.**
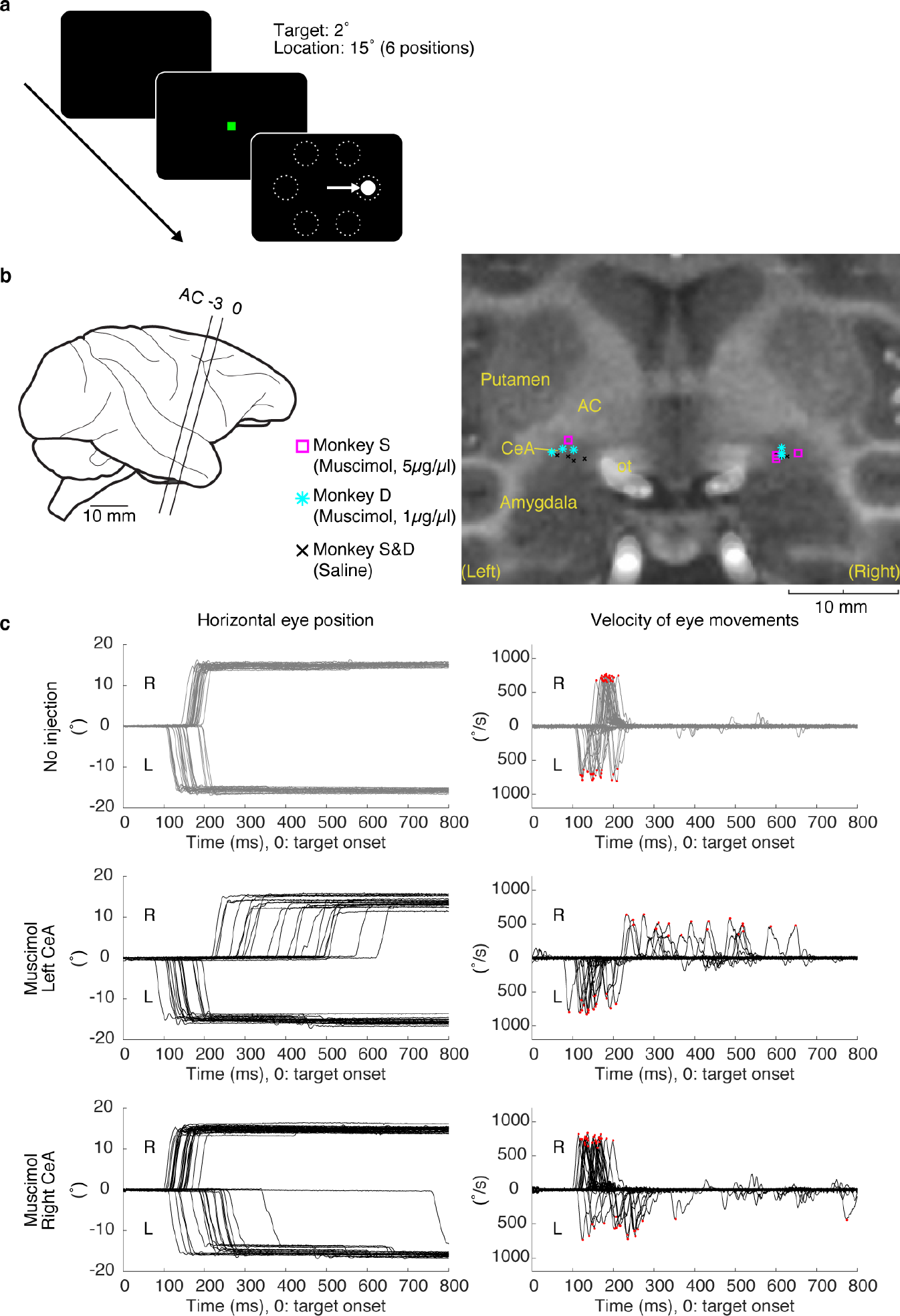
CeA inactivation suppressed saccadic eye movements to the contralateral side. **(a)** Visually-guided saccade task. **(b)** Injection sites of muscimol (GABA_A_ agonist) and saline in the left and right CeA (see Supplementary Table 1). Left: Injectrode was inserted between the two lines (15 degrees tilted anteriorly). Right: MR image along the injectrode insertion. AC: anterior commissure (e.g., −3: 3 mm posterior to AC), ot: optic tract. **(c)** Horizontal eye position (left) and velocity (right) after target onset during the visually-guided saccade task. Trials with oblique target positions are not included. Red dots show timings of the peak velocity in each saccade. Saccade latency increased and saccade velocity decreased for contralateral saccades during CeA inactivation by muscimol.

Before each injection, we identified CeA physiologically by recording neuronal activity using an injectrode (polyimide tube attached to a recording electrode for drug delivery, see Methods). We carried out 8 unilateral injections in monkey S (44 nmol muscimol: 4 times, saline: 4 times) and 12 unilateral injections in monkey D (8.8 nmol muscimol: 6 times, saline: 6 times, see Supplementary Table 1). Fig.1b shows the estimated injection sites in a magnetic resonance (MR) image (right) aligned on the orientation of the injectrode (left).

In the visually-guided saccade task, unilateral injections of muscimol into CeA suppressed contralateral saccades. Examples from a typical session are shown in Fig. 1c: left CeA inactivation (Fig. 1c, middle) increased the latencies of saccades to the right (contralateral) target compared with control data (Fig. 1c, top). Moreover, the saccade peak velocity decreased during such delayed saccades (Fig. 1c, middle right). In contrast, the saccade velocity to the left (ipsilateral) target were unchanged, although latencies were slightly shortened. Right CeA inactivation (Fig. 1c, bottom) caused the same effects: longer latencies and lower peak velocities for saccades to the left (contralateral) target and slightly shorter latencies for saccades to the right (ipsilateral) target.

We repeated the same experiment with varying delays after muscimol injections (Fig. 2a).

**Fig. 2.**
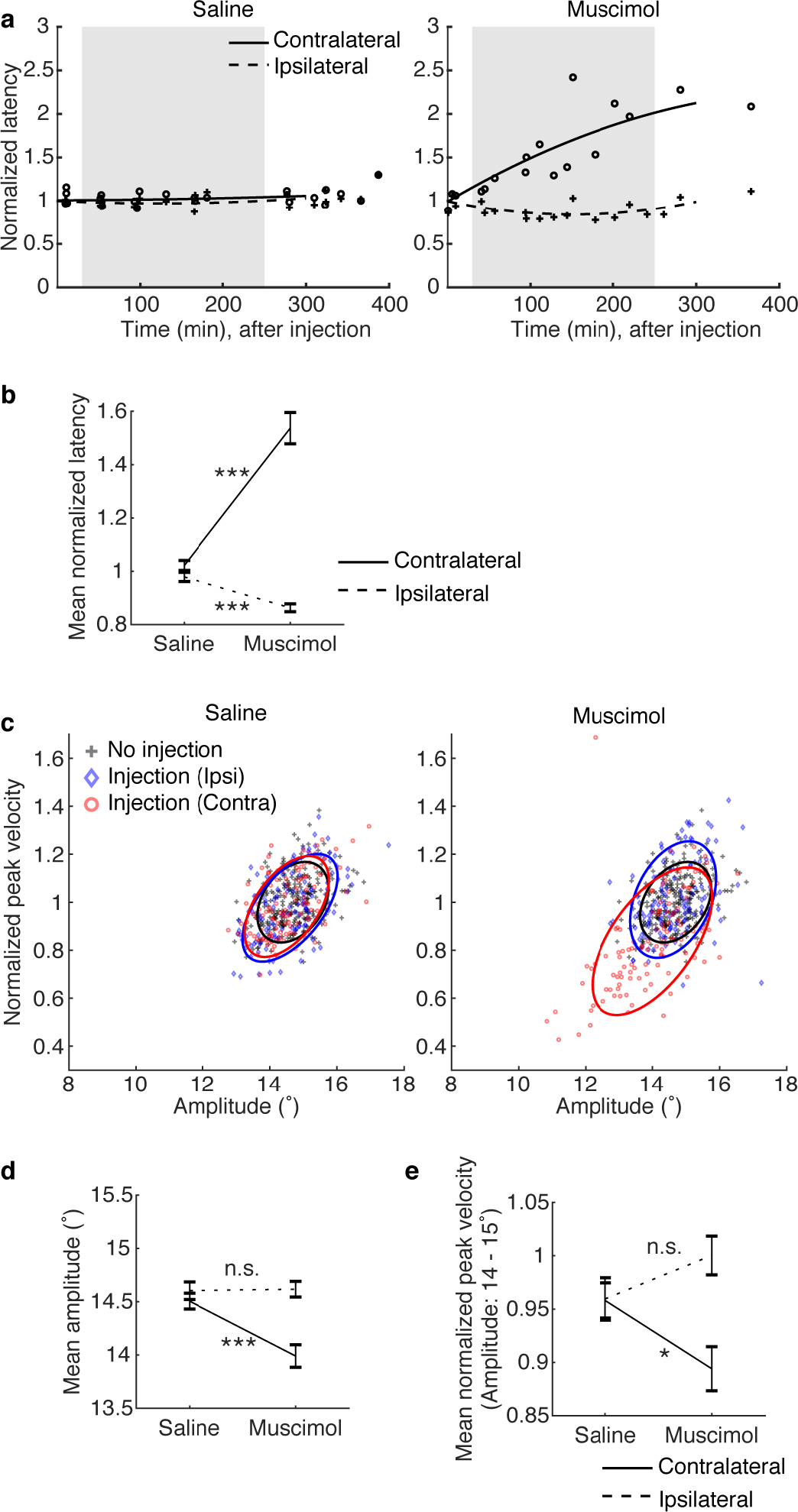
Changes in saccade features by CeA inactivation. **(a)** Change in saccade latency after injection of saline (left) and muscimol (right). Data are shown separately for saccades directed contralateral (circle) and ipsilateral (cross) to the injection site. Each data point indicates the averaged saccade latency in each session as a function of time after injection. Solid and dashed lines indicate second degree least-squares fits to the data points for contralateral and ipsilateral saccades respectively. Shaded gray area shows the effective period we used for further analysis (30-250 ms after injection). The latencies were normalized relative to those in no-injection experiments for each direction. Supplementary Fig. 3a shows non-normalized data for both monkeys. **(b)** Different effects of CeA inactivation for contralateral and ipsilateral saccades. Saccade latencies in the effective period were averaged. **(c)** Relationship between saccade amplitude (abscissa) and peak velocity (ordinate). Solid lines show 70 % confidence ellipse in each condition (black: no injection, red: CeA inactivation - contralateral saccades, blue: CeA inactivation - ipsilateral saccades). Pearson’s correlation tests show positive correlation in all conditions (monkey S: r = 0.61097, P = 1.4752e-11 in saline/ipsilateral, r = 0.5788 P = 4.3056e-10 in saline/contralateral, r = 0.36971, P = 2.3793e-05 in muscimol/ipsilateral, r = 0.59214, P = 2.0424e-12 in muscimol/contralateral; monkey D: r = 0.2288, P = 0.019484 in saline/ipsilateral, r = 0.40704 P = 2.6377e-05 in saline/contralateral, r = 0.12575, P = 0.27909 in muscimol/ipsilateral, r = 0.63926, P = 5.9775e-11 in muscimol/contralateral). **(d** & **e)** Averaged saccade amplitude and averaged peak velocity in the effective period, respectively. Note that the averaged peak velocity was calculated for saccades with amplitudes within the normal range (14-15 degrees). For (b), (d), and (e), error bars show SE. Asterisks (*) and (***) indicate statistically significant contrasts at P < 0.05 and P < 0.001. All data are for monkey S (See Supplementary Fig. 1 for monkey D).

The effect of muscimol on latency started immediately and increased over time within a daily session (Fig. 2a). We analyzed the data in an effective period (30-250 min after injection) and compared with the effect in the same window in control experiments (saline injection). Saccade latency increased significantly on the contralateral side during CeA inactivation, and the results were very similar in both monkeys (Fig. 2b and Supplementary Fig. 1b, t-test, monkey S: T(213) = −7.7457, P = 3.8129e-13, monkey D: T(182) = −9.1179, P = 1.4091e-16). On the ipsilateral side, the latency tended to decrease during CeA inactivation (monkey S: T(206.5877) = 5.0631, P = 9.1105e-07, monkey D: T(178) = 1.4341, P = 0.1533).

In addition, the amplitude of contralateral saccades sometimes became smaller (i.e., hypometric) following muscimol injection (Fig. 1c, middle and bottom) compared with control data (Fig. 1c, top). This is shown clearly in Fig. 2c-d (t-test, monkey S: T(233) = 4.0059, P = 8.3125e-05 in contralateral, T(237.7891) = −0.12437, P = 0.90113 in ipsilateral, monkey D: T(211) = 5.4291, P = 1.5507e-07 in contralateral, T(150.5153) = 0.071252, P = 0.94329 in ipsilateral). We considered the possibility that the lower peak velocity of contralateral saccades might simply be correlated with lower saccade amplitudes according to the main sequence (Pearson’s correlation, r > 0, p < 0.05) (Fig. 2c), a well-known property of saccades (ROBINSON, 1964; Baloh et al., 1975). To test this possibility, we compared the peak velocity within a short range of saccade amplitude (14-15 degree) and found that the peak velocity still significantly decreased following inactivation CeA in both monkeys (Fig. 2e, t-test, monkey S: T(80.7794) = 2.425, P = 0.017538 in contralateral, T(89.451) = −1.5122, P = 0.13402 in ipsilateral, monkey D: T(24.4669) = 4.0074, P = 0.00050223 in contralateral, T(59.5088) = 0.064379, P = 0.94888 in ipsilateral). These data indicate that inactivation of CeA alters three properties of contralateral saccades: 1) increase in latency; 2) decrease in amplitude; and 3) decrease in peak velocity.

Inactivation of CeA likewise disrupted the monkeys’ goal directed visual behavior in other phases of the task. On each trial, the monkey was required to maintain fixation on a central dot (770 ms) in order for the target object to appear (otherwise the trial ended with no opportunity for reward). Adequate fixation was disrupted by CeA inactivation in both monkeys (Supplementary Fig. 2a, Chi-square test, monkey S: P < 0.05, monkey D: P < 0.05), as they broke fixation on the central dot before the target appeared. This result, together with the preceding result (Fig. 1-2), suggests that CeA contributes to the goal-directed behavior both by suppressing inappropriate saccades in the absence of the target and facilitating appropriate saccades after target onset.

Finally, after muscimol injection into CeA, the monkeys were more prone to quit performing the behavioral tasks we tested, including the visually-guided saccade task (Supplementary Fig. 2b, mean stopping time (saline vs. muscimol): 230.5 min. vs. 120.7 min. after injection, two-sample t-test, T(14.1983) = 2.2544t, P = 0.04047). This result suggests that the role of CeA in goal-directed behavior is motivational in nature. We consider this possibility in the next section.

### Gaze shift to the ipsilateral side by CeA inactivation

In real life eye movements are used to look at things sequentially according to the animal’s attentional and motivational state (Hoffman and Subramaniam, 1995; Corbetta et al., 1998). To test whether CeA contributes to such behavior we let monkeys watch a video clip freely without any reward (Free-viewing task, duration: 5 minutes, see Methods and Fig. 3a).

**Fig. 3.**
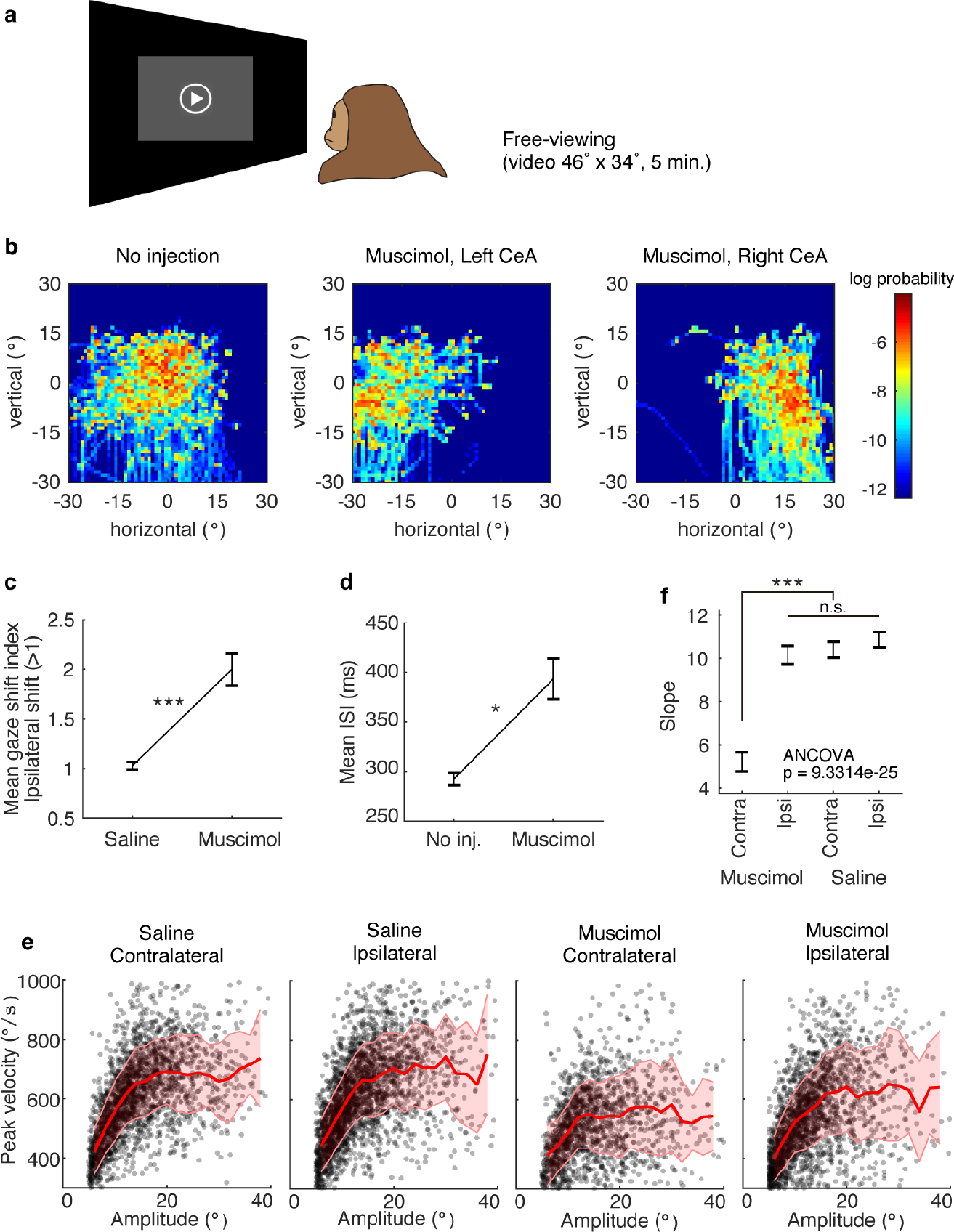
Changes in free viewing by CeA inactivation. Gaze position, saccade velocity, and amplitude during movie free-viewing. **(a)** Free-viewing task. **(b)** Gaze positions during a video clip (5 min) in three conditions in monkey S. The distribution probabilities over time after stimulus onset were calculated by logarithmic two dimensional histograms of vertical and horizontal gaze directions. **(c)** Gaze shift after saline or muscimol injection in CeA in monkey S. The shift index was calculated in each session (see Methods) and the average is plotted. **(d)** Inter-saccade interval (ISI) after no injection or muscimol injection in CeA in monkey D. **(e)** Relationship between saccade amplitude (abscissa) and peak velocity (ordinate) in monkey D. Solid lines show the average of peak velocity in each 2-degree bin of saccade amplitude. Shading shows SD. Pearson’s correlation (saline/contralateral: r = 0.51852, P = 9.6159e-155, saline/ipsilateral: r = 0.5139, P = 1.43e-169, muscimol/contralateral: r = 0.29593, P = 7.7209e-32, muscimol/ipsilateral: r = 0.46326, P = 1.9951e-107) **(f)** Slopes of regression lines (ANCOVA and post hoc, least significant difference) for the four groups shown in (e). For (c), (d), and (f), error bars show SE. Asterisks (*) and (***) indicate statistically significant contrasts at P < 0.05 and P < 0.001. The equivalent data from another subject are shown in Supplementary Fig. 4.

Each image in Fig. 3b shows eye positions of one monkey (S) during the 5-min free-viewing task. The duration of gaze (i.e., fixation between saccades) is indicated by color. Without inactivation (Fig. 3b left), monkey’s gaze was distributed in various positions, but was more common around the center. In contrast, during left or right CeA inactivation (Fig. 3b, center or right), gaze was mostly on the left (ipsilateral) or right (ipsilateral) side, respectively. These results may be characterized as “contralateral hemineglect” (Behrmann et al., 1997).

We repeated the same experiment with different timing after muscimol injection and quantified the gaze shift during the muscimol-effective period (30-250 min after injection) (Fig. 3c, see Methods). The gaze position was shifted significantly to the ipsilateral side after CeA inactivation (compared with the control data) in both monkeys (Fig. 3c and Supplementary Fig. 4c, monkey D: T(31) = −2.6607, P = 0.012238, monkey S: T(16) = −5.1959, P = 8.8354e-05).

**Fig. 4.**
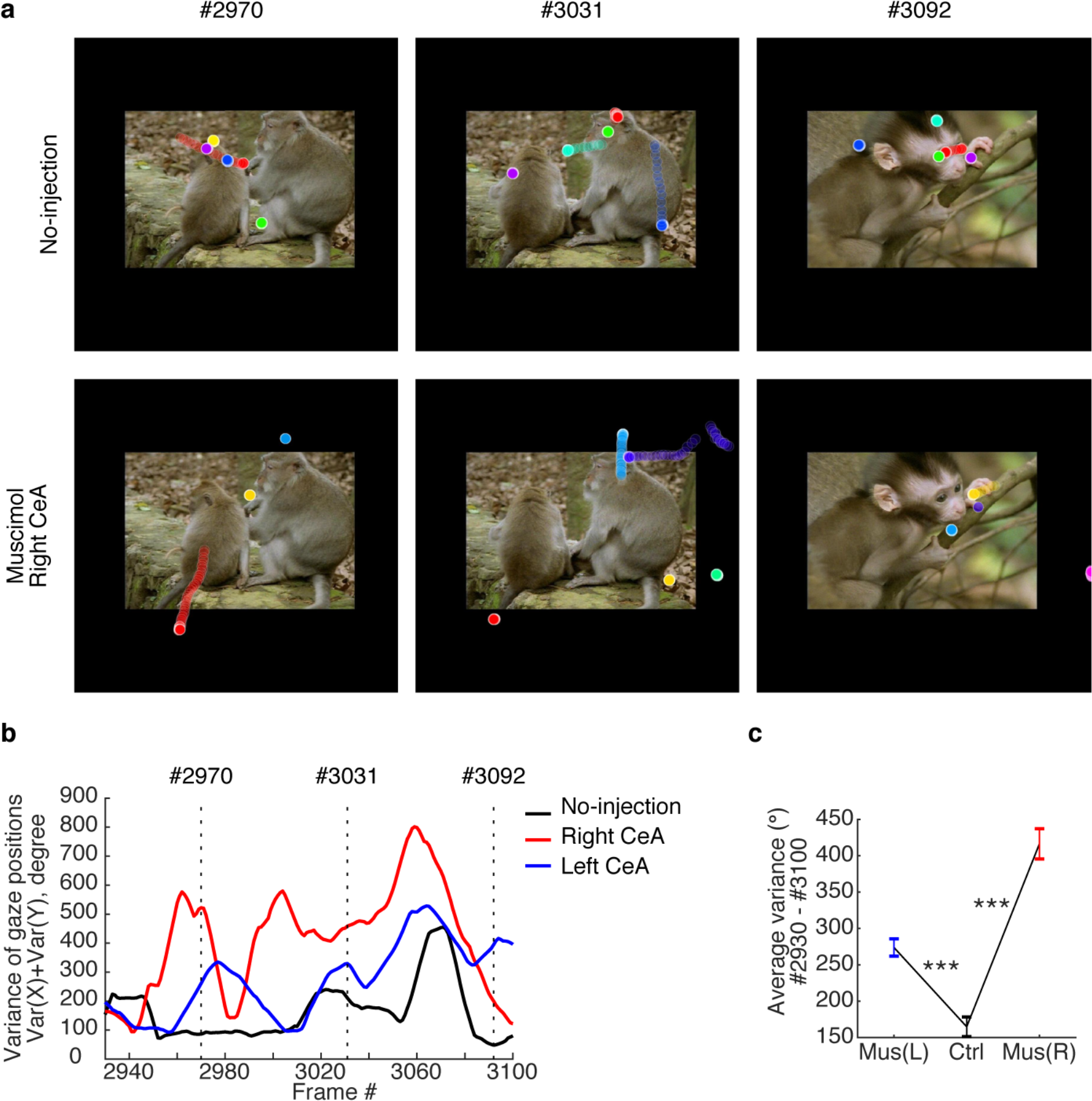
Gaze in social contexts affected by CeA inactivation. **(a)** Eye positions in selected frames of the movie in two conditions: no injection (top) and after muscimol injection in the right CeA (bottom). Different colored circles indicate eye positions in different sessions. **(b)** Across-session variance of gaze positions during movie free-viewing in three conditions. **(c)** The variance of gaze positions was increased by CeA inactivation. The data were averaged in whole sessions for each condition. Error bars show SE and asterisks (***) indicate statistically significant contrasts at P < 0.001 (one-way ANOVA, post hoc, least significant difference). All data are for monkey D (see Supplementary Fig. 5 for monkey S).

CeA inactivation also had general effects on saccadic eye movements (Fig. 3d-f). First, the frequency of saccades decreased, as the inter-saccade-interval (ISI) increased (Fig. 3d and Supplementary Fig. 4d, monkey D: T(26) = −2.7169, P = 0.011566, monkey S: T(25) = −3.177, P = 0.0039308). These results indicated that saccadic eye movements were generally suppressed, while gaze tended to stay on the ipsilateral side. This effect may be related to attention and motivation (see Discussion). Second, saccades became slower. Since the saccade amplitude varied during free viewing, the relationship between the saccade velocity and the amplitude was well estimated in each of four conditions (Fig. 3e). The peak velocity was positively correlated with the amplitude in all conditions (Pearson’s correlation, r > 0, P < 0.001). However, the slope of the regression line decreased significantly only in contralateral saccades during CeA inactivation (Fig. 3f and Supplementary Fig. 4f, ANCOVA, monkey D: P = 9.3314e-25, monkey S: P = 0.0002247). These results indicated that the saccade peak velocity decreased across different amplitudes only in saccades directed to the side contralateral to the CeA inactivation, which also confirmed the conclusion based on explicitly prompted saccades to targets in the visually-guided saccade task (Fig. 2c).

Thus far we assessed saccadic behavior over the whole five minutes duration the movie, which contained a diverse range of visual content. We next asked whether CeA inactivation altered the monkeys’ attentional and motivational state by analyzing behavior in scenes that might be of particular interest to the animals. Examples are shown in Fig. 4 for monkey D (frame #2930 - #3100) and Supplementary Fig. 5 for monkey S (frame #1210 - #1280).

**Fig. 5.**
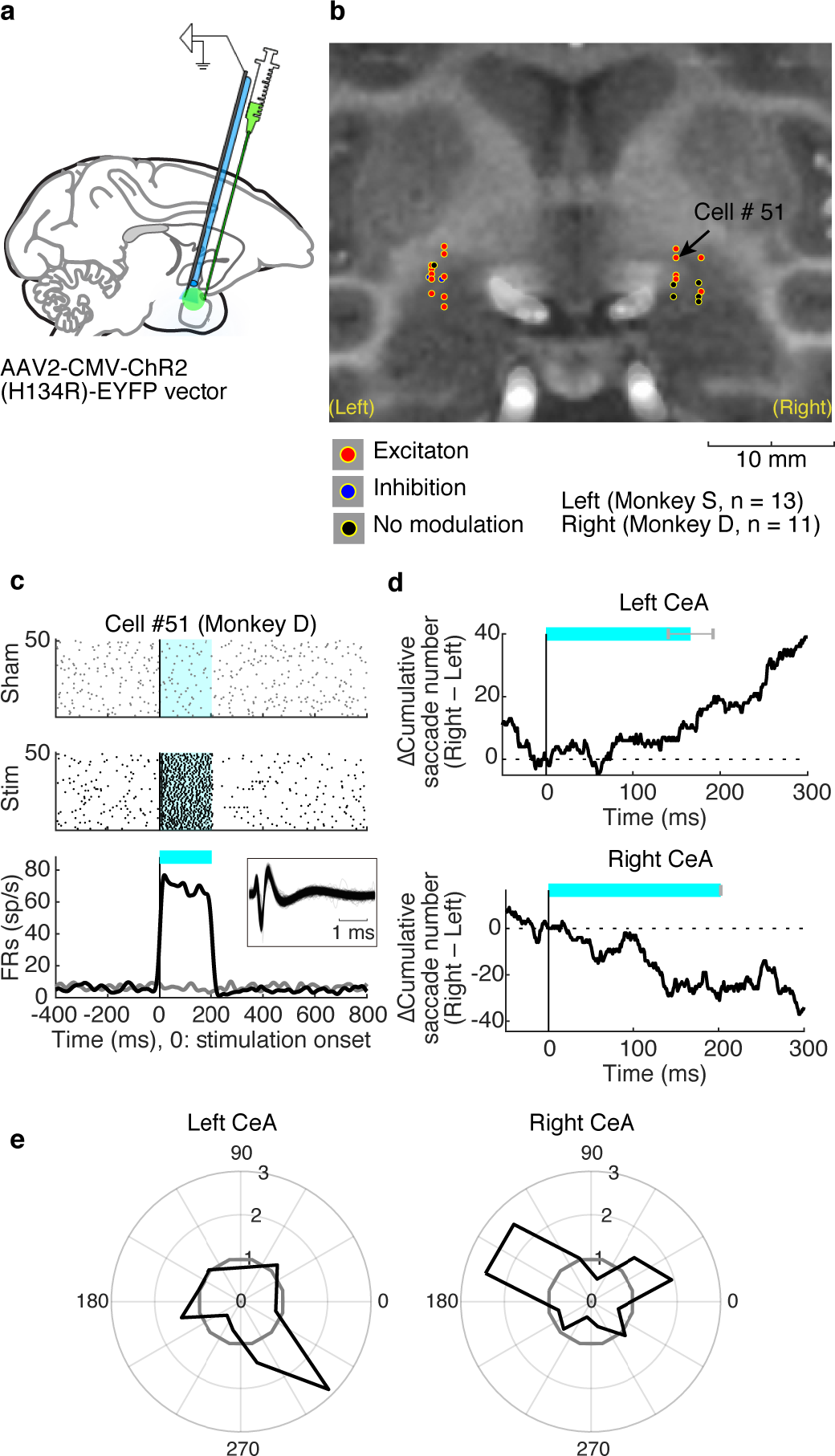
Optogenetic stimulation of CeA facilitates contralateral saccades. **(a)** Site of viral vector injection, followed by optrode placement. **(b)** Stimulation and recording positions. **(c)** Activity of an example CeA neuron that was excited during the optical stimulation period. Stimulation and non-stimulation periods were pseudo-randomly interleaved 100 times while the monkey was freely viewing during multiple visual stimulation regimes presented at random, including pictures, a blank screen, and the videos showed in the free-viewing task. Histograms and rasters are aligned at the onset of each stimulation/non-stimulation period. Spike activity was smoothed with a Gaussian kernel (σ = 10 ms). Cyan areas indicate the period of laser light emission. Inserted figure shows spike shape of the neuron. **(d)** Directional bias of saccades after CeA optical stimulation: more rightward saccades by left CeA stimulation in monkey S (top); more leftward saccades by right CeA stimulation in monkey D (bottom). Cumulative direction bias is aligned on the onset of optical stimulation (see Methods). The duration of optical stimulation somewhat varied, as shown by cyan areas and error bars (SE). **(e)** Directions of saccades that increased (>1) and decreased (<1) after CeA optical stimulation: left CeA stimulation in monkey S (left); right CeA stimulation in monkey D (right). In each direction (every 30 deg) is shown the ratio of saccades with stimulation (+) to saccades with stimulation (-).

In addition, see Supplementary Video 1 and 2 for whole images and Supplementary Fig. 6 for different periods. Gaze positions in each session are shown in a different color. Gaze was concentrated on monkey’s face and body regions in the normal condition (Fig. 4a, top). During CeA inactivation, however, gaze was often away from these regions and was scattered peripherally (Fig. 4a, bottom). The variance of gaze position was calculated in each frame (Var[horizontal] + Var[vertical], Fig. 4b). The averaged variances were significantly higher in CeA inactivation than control in both monkeys (monkey D: one-way ANOVA, F(2,510) = 62.842, P = 4.0167e-25, post hoc: least significant difference, P < 0.001; monkey S: one-way ANOVA, F(2,210) = 179.672, P = 3.3009e-46, post hoc: least significant difference, P < 0.001). The results suggest that CeA inactivation causes attention/motivation deficits to salient visual features.

### Optogenetic stimulation enhanced contralateral saccade

To further investigate the role of CeA in controlling saccadic eye movements, we stimulated CeA using optogenetic technique. This approach has the advantage of altering neuronal activity on a millisecond timescale (in contrast to the delayed effect of muscimol inactivation) (Cavanaugh et al., 2012), and further allows for simultaneous photostimulation and recording using optrodes (optic fiber attached to a recording electrode). We injected an adeno-associated virus type 2 vector (AAV2-CMV-ChR2-EYFP) into CeA of one hemisphere in both monkeys (monkey S: left, monkey D: right, Fig. 5a, see Supplementary Table 2).

After opsin expression was complete, we optically stimulated the same area randomly using 473-nm blue laser light and simultaneously recorded single-unit activity (monkey S, n = 13; monkey D, n = 11 neurons, Fig 5b) while the monkey was freely viewing. Most of CeA neurons exhibited an excitatory response during laser light emission (16 of 24, two-sample t-test P<0.05, Fig.5c), indicating that the excitatory opsin was successfully expressed in this area.

Moreover, the optical stimulation of CeA induced contralateral saccades (Fig. 5d-e). Monkey S made saccades to the right side more frequently during and after the optical stimulation in the left CeA (Fig. 5d top and 5e left), whereas monkey D made those to the left side upon the optical stimulation in the right CeA (Fig. 5d bottom and 5e right). The cumulative number of saccades in each session was significantly greater on the contralateral side, compared with no stimulation control (right-tailed one-sample t-test, T(23) = 2.6434, P = 0.0072624). These results further confirmed that CeA facilitates saccadic eye movements to the contralateral side.

## Discussion

We have found that the amygdala facilitates saccadic eye movements in a highly lateralized manner. Our recent study showed that many neurons in CeA were sensitive to emotional and environmental contexts (i.e., dangerous, rich, poor), and their activity was tightly correlated with reaction times of goal-directed saccades across contexts (Maeda et al., 2018). These findings suggest that the amygdala facilitates goal-directed behavior specifically in emotionally charged contexts. The current study supports this hypothesis, as the inactivation of CeA suppressed saccades by increasing the latency and decreasing the amplitude and velocity (Fig. 1 and 2). Notably, these effects were restricted to objects in the visual field contralateral to the inactivated CeA. This spatial selectivity of CeA control of eye movements was manifest from three independent sources of evidence. First, the suppression occurred only in saccades directed to the contralateral side (Fig. 1 and 2). Second, during free-viewing, gaze position was shifted to the ipsilateral side (i.e., away from the contralateral side) (Fig. 3). Finally, optogenetic stimulation of CeA enhanced saccades to the contralateral side (Fig. 5).

Taken together, these findings point to a role for the amygdala as an oculomotor control structure - a conclusion that may seem surprising, since (with the exception of a few studies (Janak and Tye, 2015; Han et al., 2017)) the amygdala is generally understood to subserve emotional rather than motor function.

The idea of an oculomotor amygdala may be brought into harmony with the prevailing view by appreciating the fact that eye movement deficits reported here were likely associated with emotion, motivation, and social attention. Specifically, CeA inactivation caused the following deficits in motivation and/or social attention: 1) reduced the probability of waiting for the target in each trial (Supplementary Fig. 2a); 2) shortened the duration of task performance within a day (Supplementary Fig. 2b); 3) reduced the frequency of saccades during movie free-viewing (Fig. 3d); and 4) reduced the likelihood of gaze towards social interaction during movie free-viewing (Fig. 4, Supplementary Fig. 5, and Supplementary Fig. 6). These results are consistent with the findings of previous studies (Emery, 2000; Mosher et al., 2014; Dal Monte et al., 2015; Forcelli et al., 2016; Minxha et al., 2017; Taubert et al., 2018).

Together with our previous findings (Maeda et al., 2018), these results suggest that CeA governs social and emotional behavior on multiple levels. At the most general level, CeA mediates context-appropriate social orienting, as evidenced by the effect of CeA inactivation during movie free-viewing. At a more fine-grained level, CeA also selectively mediates the spatially-oriented action for finding objects of interest quickly and accurately. This goal-directed aspect of amygdala function was disrupted by CeA inactivation and facilitated by CeA optogenetic stimulation during the simple saccade task. Both levels must work in tandem for appropriate behavior to take place, as the general promotion of a context-appropriate behavioral regime must be followed up by the narrowly targeted actions of choosing or rejecting particular objects. To this end, the behavior must be controlled accurately, especially in the spatial domain, otherwise random involuntary movement would prevail. In this sense, the spatially selective (i.e., contralateral) control of saccades by CeA is critical.

What is the neuronal mechanism of saccadic control by CeA? Anatomically, CeA projects to several structures that contribute to saccadic eye movements, including SNr, the globus pallidus, the striatum, the zona incerta, and the reticular formation (Price and Amaral, 1981; Shinonaga et al., 1992; Fudge et al., 2002; Fudge et al., 2004; Cho et al., 2013; Griggs et al., 2017; Amita et al., 2018). Importantly, most output neurons in CeA are GABAergic and inhibitory (Tovote et al., 2015), and saccades are controlled mainly by SC (Hikosaka et al., 2000). Thus, the activation of CeA output should lead to a disinhibition of SC. This scheme suggests the following circuit mechanism: CeA neurons have direct inhibitory connections to SNr (Lee et al., 2010), which in turn has inhibitory connections to SC (Hikosaka and Wurtz, 1983). Because the firing rates of many CeA neurons increases in contexts that arouse animals (e.g., dangerous or rich), the final effect would be to facilitate saccades specifically in exciting contexts (Maeda et al., 2018). In summary, our results reveal that the amygdala plays an important, previously unappreciated role in the control of saccadic eye movements.

## Methods

### Subjects and surgery

We used four hemispheres of two rhesus monkeys (Macaca mulatta) in this study (monkey S: 8.5 kg, 7y old, male, monkey D: 5.0 kg, 13y old, female). All animal care and experimental procedures were approved by the National Eye Institute Animal Care and Use Committee and complied with the Public Health Service Policy on the Humane Care and Use of Laboratory Animals. Both animals underwent surgery under general anesthesia during which a head holder and a recording chamber were implanted on the head. Based on a stereotaxic atlas (Saleem and Logothetis, 2007), we implanted a rectangular chamber targeting the amygdala. The chamber was tilted anteriorly by 15 degrees in both monkeys. After confirming the position of recording chamber using MRI, a craniotomy was performed during a second surgery.

### Muscimol Inactivation

Head position was stabilized throughout the experimental session by means of a chronically implanted head post. After the electrophysiological recording (mapping) of amygdala neurons in both monkeys, we performed inactivation experiments to test for a causal relationship between CeA neuronal activity and eye movements. To accurately inactivate the brain structure, we used an electrode assembly (injectorode) consisting of an epoxy-coated tungsten microelectrode (FHC) for unit recording and a polyimide tube (MicroLumen) for drug delivery. After the precise identification of CeA by unit recording, we injected the GABA_A_ receptor agonist muscimol (Sigma; 8.8 nmol [1µg] or 44 nmol [5 µg] in a 1 µl volume) into the CeA in one hemisphere. Because the effect of muscimol on CeA lasted for several hours, inactivation sessions were limited to twice per week. Soon after the injection was completed (within 5 min), the animal was invited to start several tasks and repeat performance after periodic rest breaks every 20-60 min for 3-5 hr. In. this study, we used visually-guided saccade task and free-viewing task to analyze inactivation effects for eye-movements.

### Experimental control

All behavioral tasks were controlled by a custom neural-recording and behavior-controlling system (Blip; available at http://www.robilis.com/blip/). The monkey sat in a primate chair facing a fronto-parallel screen in a sound-attenuated and electrically shielded room. Visual stimuli were rear-projected on the screen by a digital light processing projector (PJD8353s, ViewSonic). Eye position was sampled at 1 kHz using a video-based eye tracker (EyeLink 1000 Plus, SR Research).

### Visually-guided saccade task

Trials began with the appearance of a central fixation point (Fig. 1a). After 770 ms of fixation the fixation point disappeared, and a saccadic target (2 × 2 degree) was presented on one of six positions randomly. The monkey was required to make a saccade and maintain fixation for 400 ms within the target window (10 × 10 degree) to obtain a water reward.

### Free-viewing task

The monkey freely watched a movie without any rewards. The movie consisted of several video clips and lasted 5 minutes (30 frames/sec), mostly showing macaque monkeys engaged in natural behavior. We used the same movie across experiments. The video clips were assembled from commercially available documentaries and wildlife footage and previously used in another study (McMahon et al., 2015). The movie was presented within a rectangular frame of 46° wide and 34° high by custom-written MATLAB functions.

### Optogenetic experiments

After the muscimol injection experiments were completed, we injected an adeno-associated virus type 2 vector (AAV2-CMV-ChR2-EYFP: 9.0 × 10^12^ genome copy/ml) into CeA area of one hemisphere in both monkeys (monkey S: left CeA, monkey D: right CeA). The vector was successfully used in the macaque brain in a previous study (Inoue et al., 2015). Two or three penetrations were made into one side of CeA at least 1.41 mm apart from each other (Supplementary Table 2]. For each penetration, 2–3 µl of the vector was introduced one time with rate 0.4 µl/min for initial 0.2 µl, then 0.08 µl/min for rest of them by using the injectrode with 10 µl Hamilton syringe and motorized infusion pump (Harvard Apparatus, Holliston, MA, USA).

For optical stimulation and electrophysiological recording, we used optrodes consisting of an epoxy-coated tungsten microelectrode (FHC) for unit recording and an optic fiber (200 µm diameter, Doric Lenses) for optical stimulation using laser light. The light sources were 473nm DPSS blue light lasers with a maximum power of up to 100mW (Opto Engine LLC). We left the laser on continuously during the experiment and placed a mechanical switcher (Luminos Industries Ltd) in the light path to turn the laser on and off. We measured the light intensity at the tip of the optrode before penetration of the brain using an optical power meter (1916-C, Newport Corporation) coupled with a 818-SL/DB photo detector. The light intensities were 30-50 mW/mm^2^ in this experiment.

### Data analysis for saccade

In the visually-guided saccade task, the saccade latency was measured as the time between the offset of the fixation point (simultaneous with the onset of the object) and the onset of the saccade to the fractal object in the visually-guided saccade task. First, saccades were detected when the peak velocity of the polar component exceeded 300 degrees/s. Saccade onset time was defined as the time point preceding the detected saccade at which the velocity exceeded 30 degrees/s. In the free-viewing task, we analyzed saccades with amplitudes ranging from 5 to 40 degrees. In the optogenetic stimulation experiments, differences in the cumulative number of saccades between stimulation and control conditions were plotted, including all sessions in each monkey. Saccadic eye movements to right and left directions were applied to positive and negative numbers, respectively. Positive number indicates increasing saccadic eye movement to right side comparing with control data and vice versa.

## Acknowledgements

We thank D. McMahon for manuscript-writing assistance; H. Amita for assistance with optogenetic experiments; and M.K. Smith, D. Parker, D. O’Brien, V.L. McLean, I. Bunea, G. Tansey, D. Yu, A.M. Nichols, T. W. Ruffner, J.W. McClurkin, J. Fuller-Deets, and A.V. Hays, and M Fujiwara for technical assistance. This work was supported by Intramural Research Program at the National Institutes of Health, National Eye Institute (project number: 1ZIAEY000415-16, https://projectreporter.nih.gov/project_info_details.cfm?aid=9796699&map=y), the Japan Agency for Medical Research and Development (Grants JP18dm0307021 and JP18dm0207003), and the Japan Science and Technology Agency (PRESTO Grant JPMJPR1683).

## Author contributions

K.M. and O.H. designed this research, performed the experiments, analyzed the data, and wrote the manuscript. K.I. and M.T. produced the viral vector. K.M., K.I., J.K., M.T., and O.H. discussed the results.

## Data Availability

The data that support the findings of this study are available from the corresponding author upon reasonable request.

## Supplementary

**Supplementary schematic.**
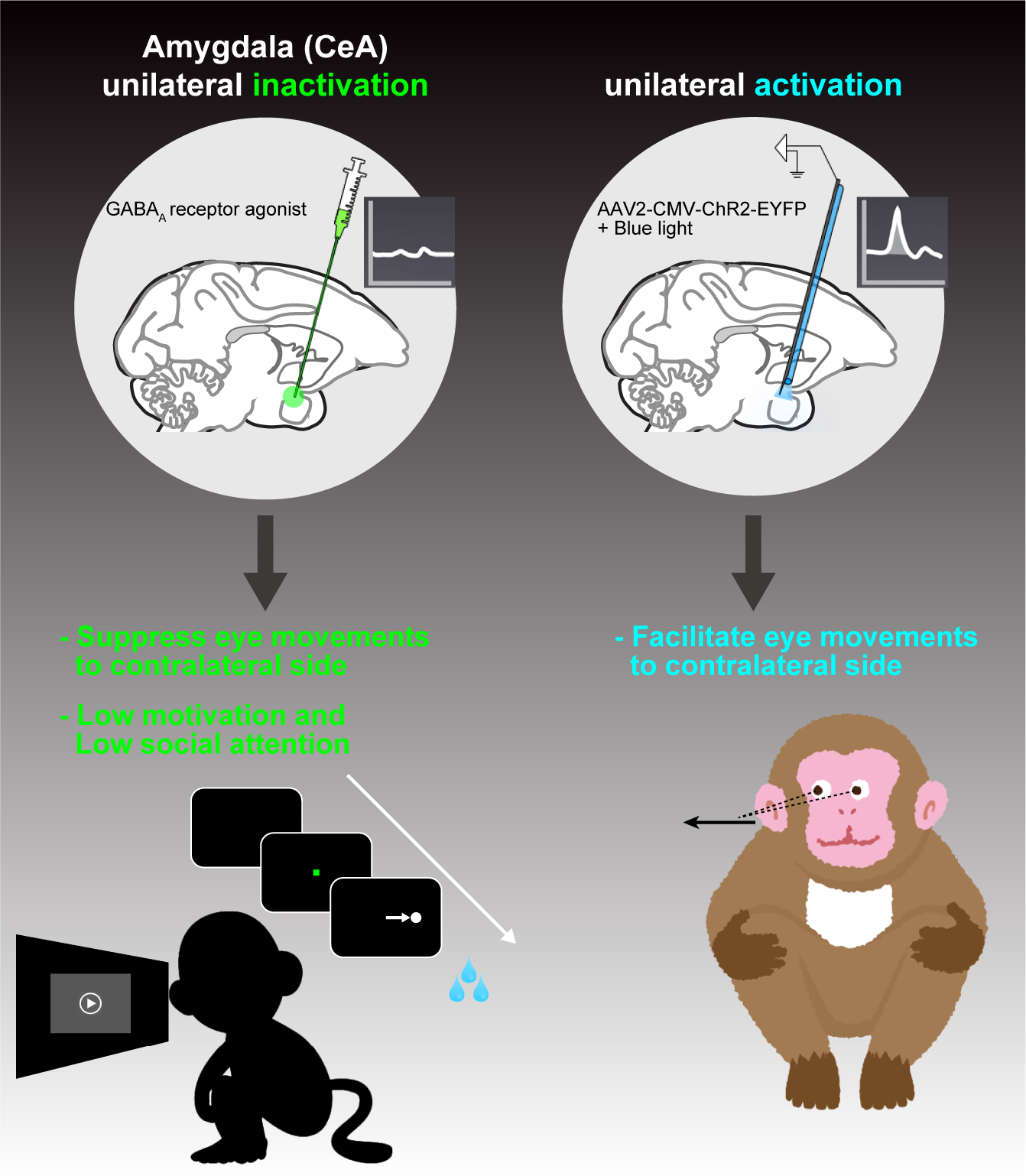

**Supplementary Fig. 1.**
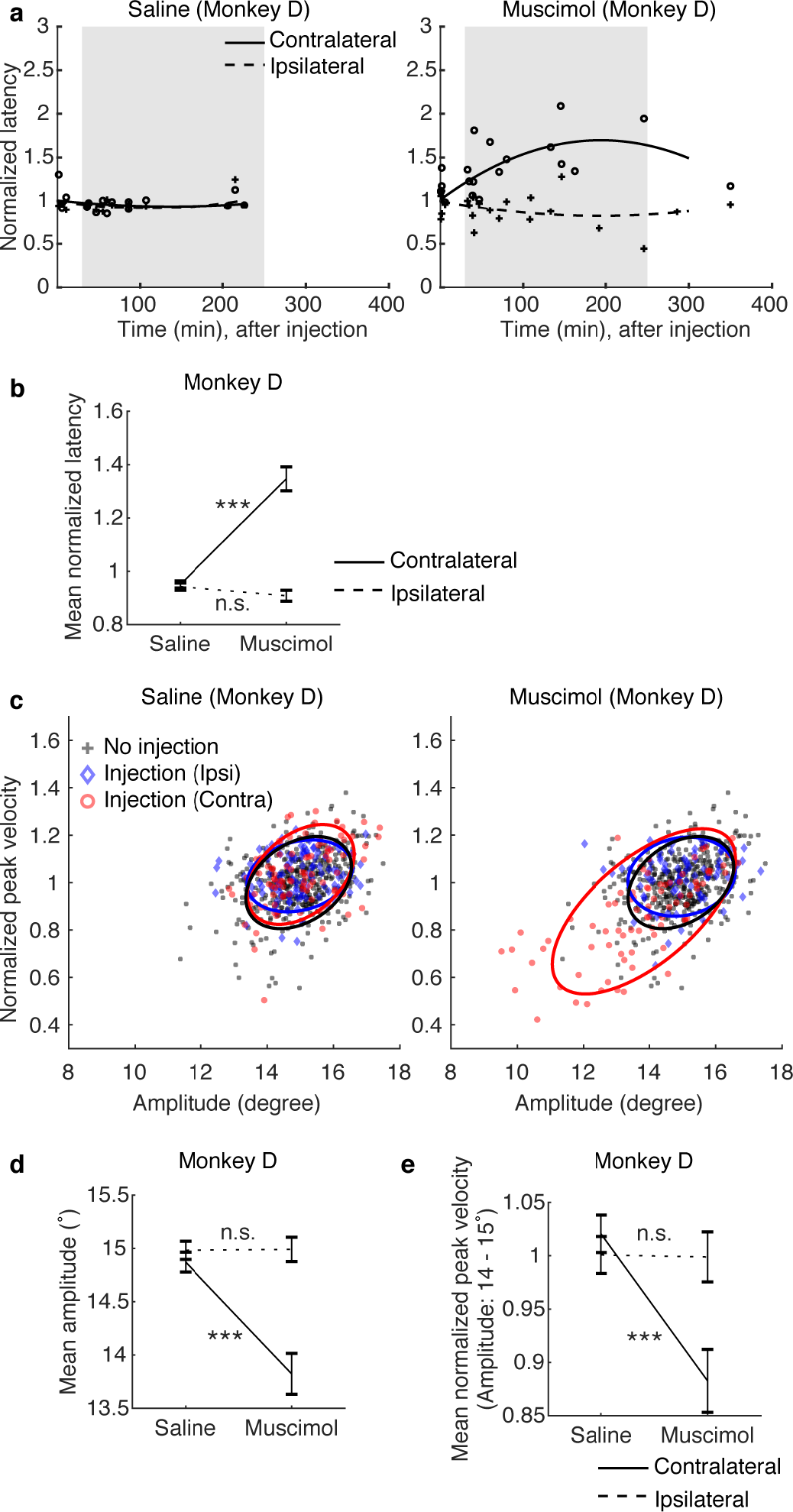
Data from another subject (related to Fig. 2). For (b), (d), and (e), error bars show SE; asterisk (***) indicates statistically significant contrasts at P < 0.001.

**Supplementary Fig. 2.**
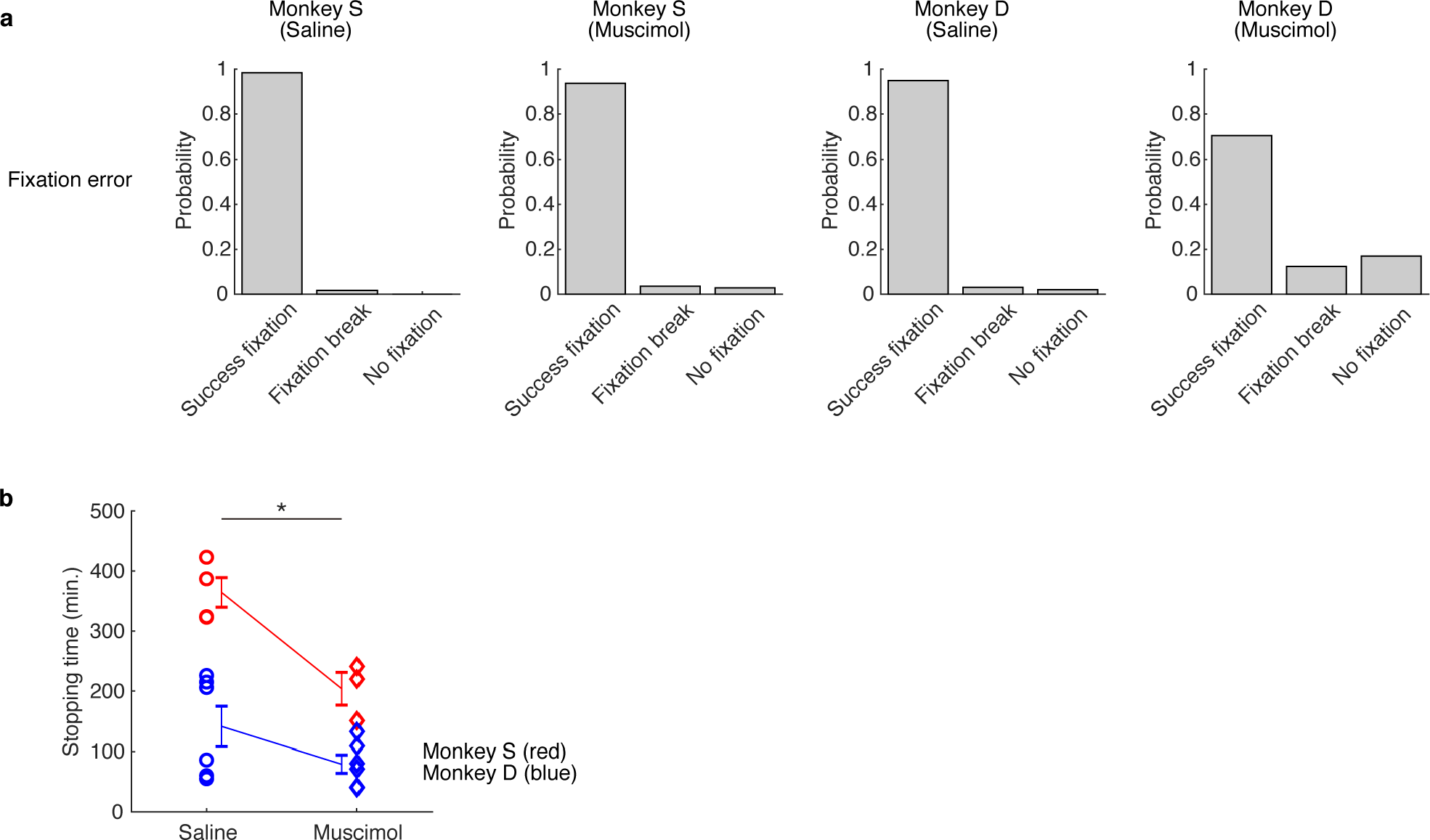
Motivational deficits during goal-directed behavior by CeA inactivation. **(a)** Increase in fixation breaks before saccade target appeared. **(b)** Earlier stopping of the visually-guided saccade task. The stopping time was recorded when the monkey made no fixation in five consecutive trials in each injection session. When there is no such error, the time of the final session was recorded as the stopping time. Each marker indicates the stopping time in each injection experiment (red: monkey S, blue: monkey D). Error bars show SE; asterisk (*) indicates statistically significant contrasts at P < 0.05.

**Supplementary Fig. 3.**
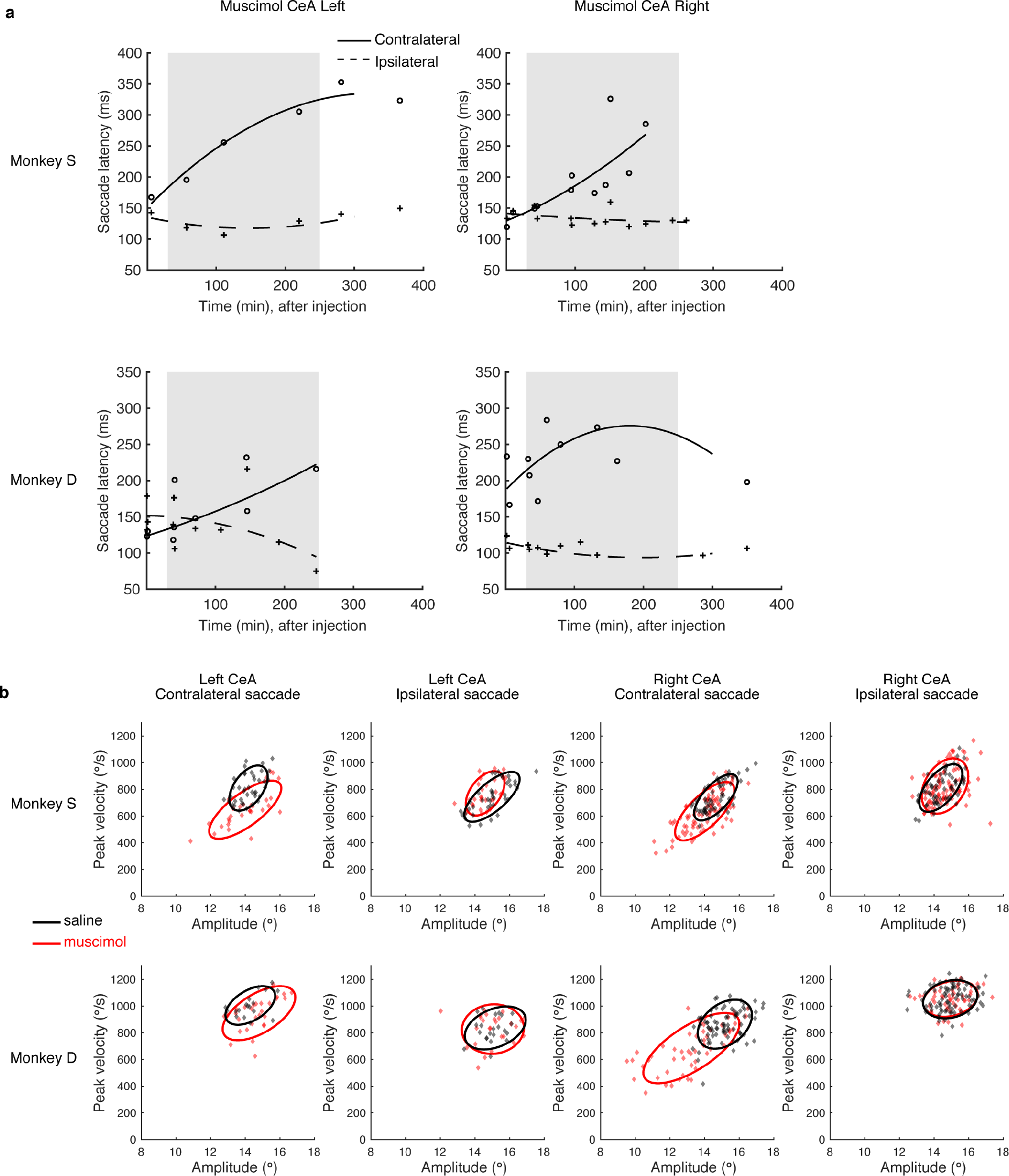
(a) Original data of Fig. 2a and Supplementary Fig. 1a. (b) Original data of Fig. 2c and Supplementary Fig. 1c.

**Supplementary Fig. 4.**
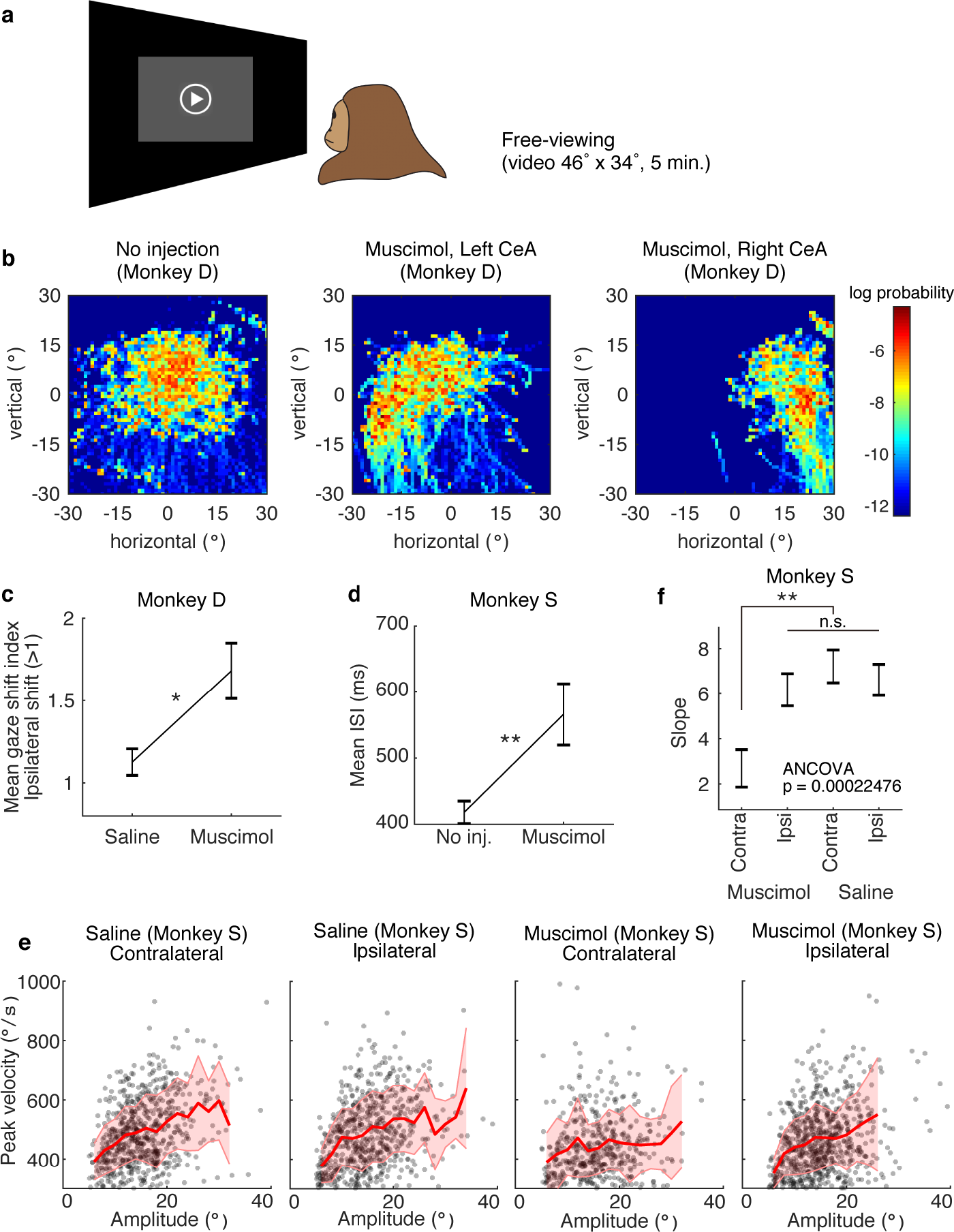
Data from another subject (related to Fig. 3). For (c), (d), and (f), error bars show SE; asterisks (*) and (**) indicate statistically significant contrasts at P < 0.05 and P < 0.01.

**Supplementary Fig. 5.**
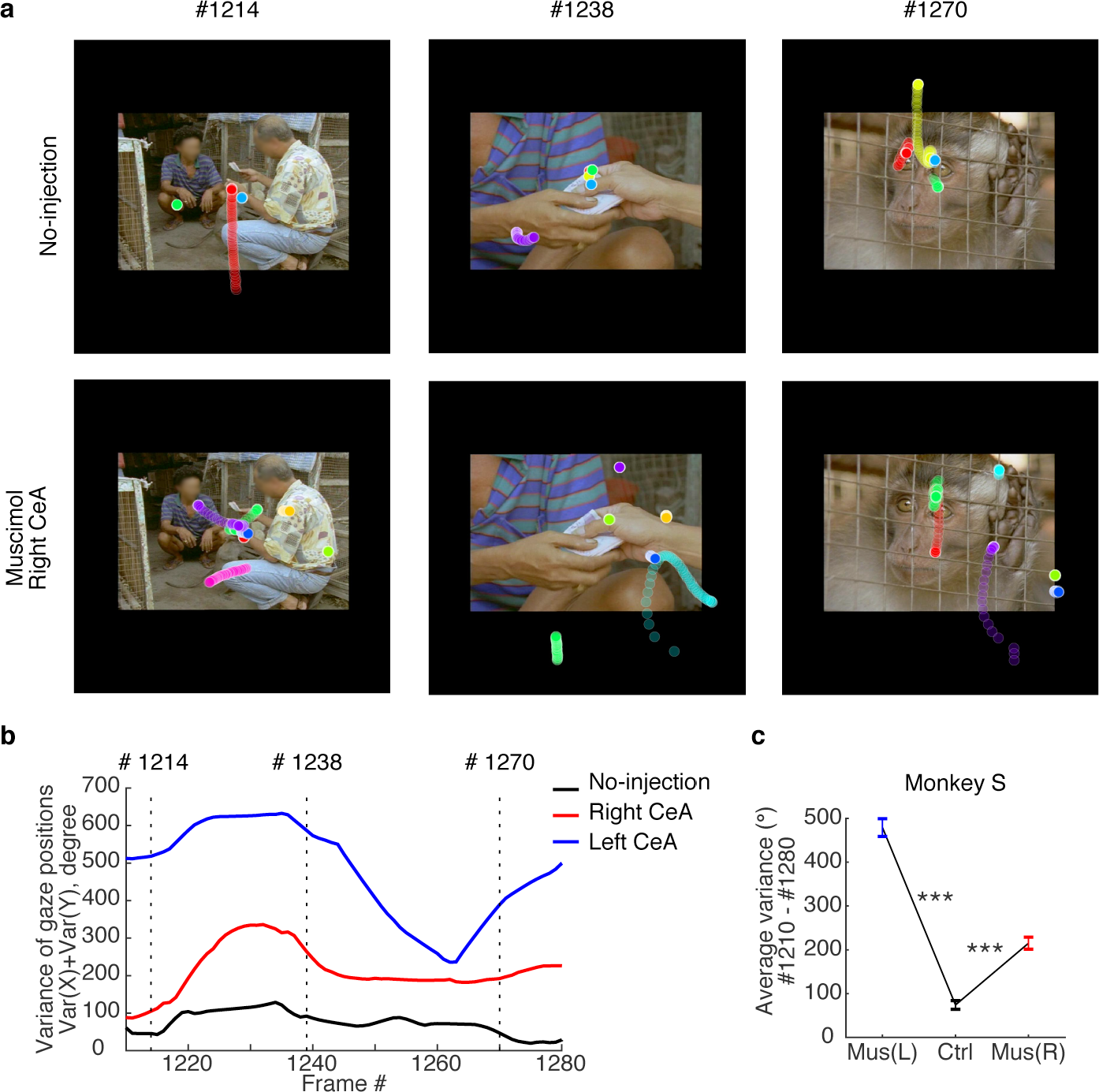
Data from another subject (related to Fig. 4). For (c), error bars show SE; asterisk (***) indicates statistically significant contrasts at P < 0.001. Face images are blurred to avoid the inclusion of identifying information of people.

**Supplementary Fig. 6.**
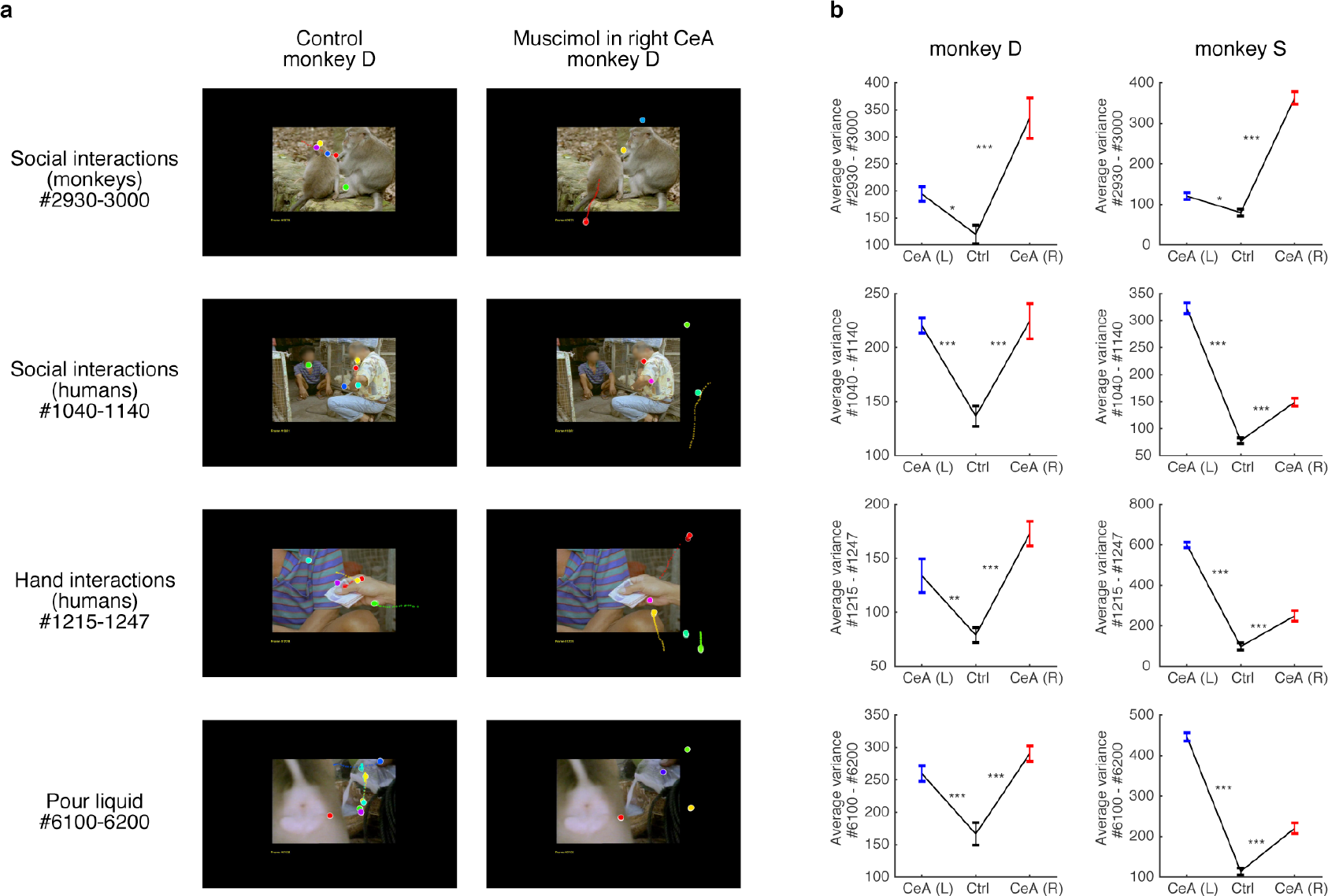
Gaze positions in different periods of video clips (related to Fig. 4 and Supplementary Fig. 5). Error bars show SE; asterisks (*), (**), and (***) indicate statistically significant contrasts at P < 0.05, P < 0.01, and P < 0.001. Face images are blurred to avoid the inclusion of identifying information of people.

**Supplementary Table 1.** Detail of muscimol injection experiments (task VS: visually-guided saccade, task FV: free-viewing task, Mus.: muscimol). AC: anterior commissure. The positions listed below are chamber coordinates, which were tilted 15 degrees anteriorly relative to the stereotaxic reference frame.

| Monkey | Concn., $\mu\text{g}/\mu\text{l}$ | Volume ($\mu\text{l}$) | Injection side | AP position from AC | Lateral from AC | Depth from AC | Tasks |
| --- | --- | --- | --- | --- | --- | --- | --- |
| D | Mus. $1\mu\text{g}/\mu\text{l}$ | 1 | Left | -2 | 9 | 5.9 | VS, FV |
| D | Mus. $1\mu\text{g}/\mu\text{l}$ | 1 | Left | -3 | 11 | 6.1 | VS, FV |
| D | Mus. $1\mu\text{g}/\mu\text{l}$ | 1 | Left | -1 | 10 | 5.8 | VS, FV |
| D | Mus. $1\mu\text{g}/\mu\text{l}$ | 1 | Right | -2 | 10 | 6.2 | VS, FV |
| D | Mus. $1\mu\text{g}/\mu\text{l}$ | 1 | Right | -2 | 10 | 5.7 | VS, FV |
| D | Mus. $1\mu\text{g}/\mu\text{l}$ | 1 | Right | -1 | 10 | 6.4 | VS, FV |
| D | Saline | 1 | Right | -2 | 10 | 6.3 | VS, FV |
| D | Saline | 1 | Right | 0 | 10 | 6.5 | VS |
| D | Saline | 1 | Right | 0 | 10 | 6.3 | VS, FV |
| D | Saline | 1 | Right | -2 | 10 | 6.5 | VS, FV |
| D | Saline | 1 | Left | -1 | 8 | 6.7 | VS, FV |
| D | Saline | 1 | Left | 0 | 9 | 6.9 | VS, FV |
| S | Mus. $5\mu\text{g}/\mu\text{l}$ | 1 | Left | -1 | 9.5 | 5 | VS, FV |
| S | Mus. $5\mu\text{g}/\mu\text{l}$ | 1 | Right | -3 | 11.5 | 6.2 | FV |
| S | Mus. $5\mu\text{g}/\mu\text{l}$ | 1 | Right | -1 | 9.5 | 6.4 | VS, FV |
| S | Mus. $5\mu\text{g}/\mu\text{l}$ | 1 | Right | -1 | 9.5 | 6.7 | VS, FV |
| S | Saline | 1 | Left | -1 | 9.5 | 6.5 | VS, FV |
| S | Saline | 1 | Right | -1 | 9.5 | 6.2 | VS, FV |
| S | Saline | 1 | Right | -2 | 10.5 | 6.5 | VS, FV |
| S | Saline | 1 | Left | -2 | 7.5 | 6.4 | VS, FV |

**Supplementary Table 2.**
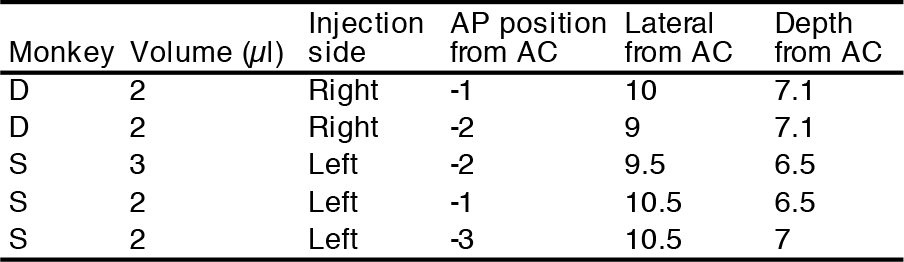
Injection sites of the viral vector for optogenetic experiments. AC: anterior commissure. The positions listed below are chamber coordinates, which were tilted 15 degrees anteriorly relative to the stereotaxic reference frame.

| Monkey | Volume (μl) | Injection side | AP position from AC | Lateral from AC | Depth from AC |
| --- | --- | --- | --- | --- | --- |
| D | 2 | Right | -1 | 10 | 7.1 |
| D | 2 | Right | -2 | 9 | 7.1 |
| S | 3 | Left | -2 | 9.5 | 6.5 |
| S | 2 | Left | -1 | 10.5 | 6.5 |
| S | 2 | Left | -3 | 10.5 | 7 |

**Supplementary Video 1**

Example eye positions during free-viewing task in normal condition (speed x0.2)

**Supplementary Video 2**

Example eye positions during free-viewing task after muscimol injection into right CeA (speed x0.2)

